# Shared somatosensory–motor neural population dynamics track motor recovery after stroke

**DOI:** 10.64898/2026.02.10.705199

**Authors:** Ian S. Heimbuch, Preeya Khanna, Lisa Novik, Robert J. Morecraft, Karunesh Ganguly

## Abstract

Although somatosensation is known to be important for precise hand control, how coordinated activity across sensory and motor cortical areas supports hand movements and object interactions is unclear. We simultaneously recorded neural activity from dorsal premotor cortex (PMd) and Area 2 of the primary somatosensory cortex in three rhesus macaques performing a reach-to-grasp task during recovery from a primary motor cortex (M1) lesion. Using dual-area latent factor analysis, we decomposed population activity into cross-area factors (CFs) shared between PMd and Area 2 and within-area factors (WFs) local to each region. Shared CFs tracked hand kinematics closely despite explaining less overall variance. Strikingly, the emergence of long-timescale, stereotyped sensory–motor cross-area dynamics was correlated with recovery of prehension and object grasping, whereas changes in within-area dynamics were not correlated with recovery. These findings suggest that restoration of coordinated sensory–motor population dynamics is important for recovery of skilled hand function after injury.

## Introduction

Sensorimotor function relies on a highly interconnected network involving multiple motor and sensory areas.^1–3^ Past work, e.g., using lesions of sensory cortex in non-human primates^4,5^ and the pharmacological numbing of fingertips in humans^6– 8^, has demonstrated that sensory inputs are very important for precise hand control. Despite this, little is known about how coordinated activity patterns across motor and sensory cortices support reach-to-grasp movements and object interactions, particularly during recovery from injury. To address this, we recorded neural activity from dorsal premotor cortex (PMd), which contributes to the planning and generation of coordinated motor actions,^9,10^ and area 2 of parietal cortex, which integrates proprioceptive and tactile signals,^11–16^ during recovery of arm and hand function following a lesion to the hand and arm region of primary motor cortex (M1). Notably, past work has demonstrated that premotor areas in general^17–20^, and PMd specifically, are sites that support recovery of upper-limb function after the loss of M1.^21–23^ Our lesion model thus provides an opportunity to assess how recovery of hand dexterity is correlated with changes in coordinated population activity across motor and sensory regions and activity local to each region. We specifically decomposed population-level activity in PMd and Area 2 into two distinct patterns: those that are explicitly shared between the two regions and those that are local to each region.^24^ Based on prior work showing that recovery after motor cortical injury engages widespread structural plasticity^25^ and can be influenced by interventions that modulate distributed activity,^21,23^ we hypothesized that signals shared across motor and somatosensory areas is more predictive of recovery of dexterity than activity confined to a single region.

To test how coordinated activity across sensory–motor areas is linked to recovery of hand dexterity, we simultaneously recorded neural spiking activity from PMd and Area 2 in three animals performing a reach-to-grasp single pellet task during recovery from the M1 lesion. Using two-area dimensionality reduction,^24^ we identified cross-area latent factors (CFs) that reflect shared PMd–Area 2 neural population dynamics. Interestingly, these CFs tracked hand kinematics more closely and predicted wrist speed more accurately than activity confined to a single region. During recovery, the time constants of shared CFs became progressively longer and there was reduced single trial variability; these were correlated with recovery of task performance. In contrast, local within-area activity changes were not significantly correlated with recovery. Together, these results suggest that the restoration of shared sensory and motor population dynamics is important for recovery of prehension after injury.

## Results

### Activity during reach-to-grasp actions in PMd and Area 2 in recovered animals

Rhesus macaques were trained to perform a reach-to-grasp task (Figure 1A) in which animals retrieved pellets from wells of varying diameters.^21^ Animals were trained until performance plateaued. Next, animals underwent stroke induction in which the arm and hand area of the right-hemisphere primary motor cortex (M1), contralateral to the trained left hand, was targeted with surface vessel occlusion followed by subpial aspiration.^26^ Two of the three animals (Monkeys Ba and H) showed significant reach-to-grasp kinematic deficits following M1 stroke (Figure 1A, Panel 3), followed by gradual behavioral recovery over the course of weeks (Figure 1A, Panel 4). In the third animal (Bk), despite a similar lesion induction, reach-to-grasp function in the first post-stroke session was not significantly different from pre-stroke levels.

**Figure 1.**
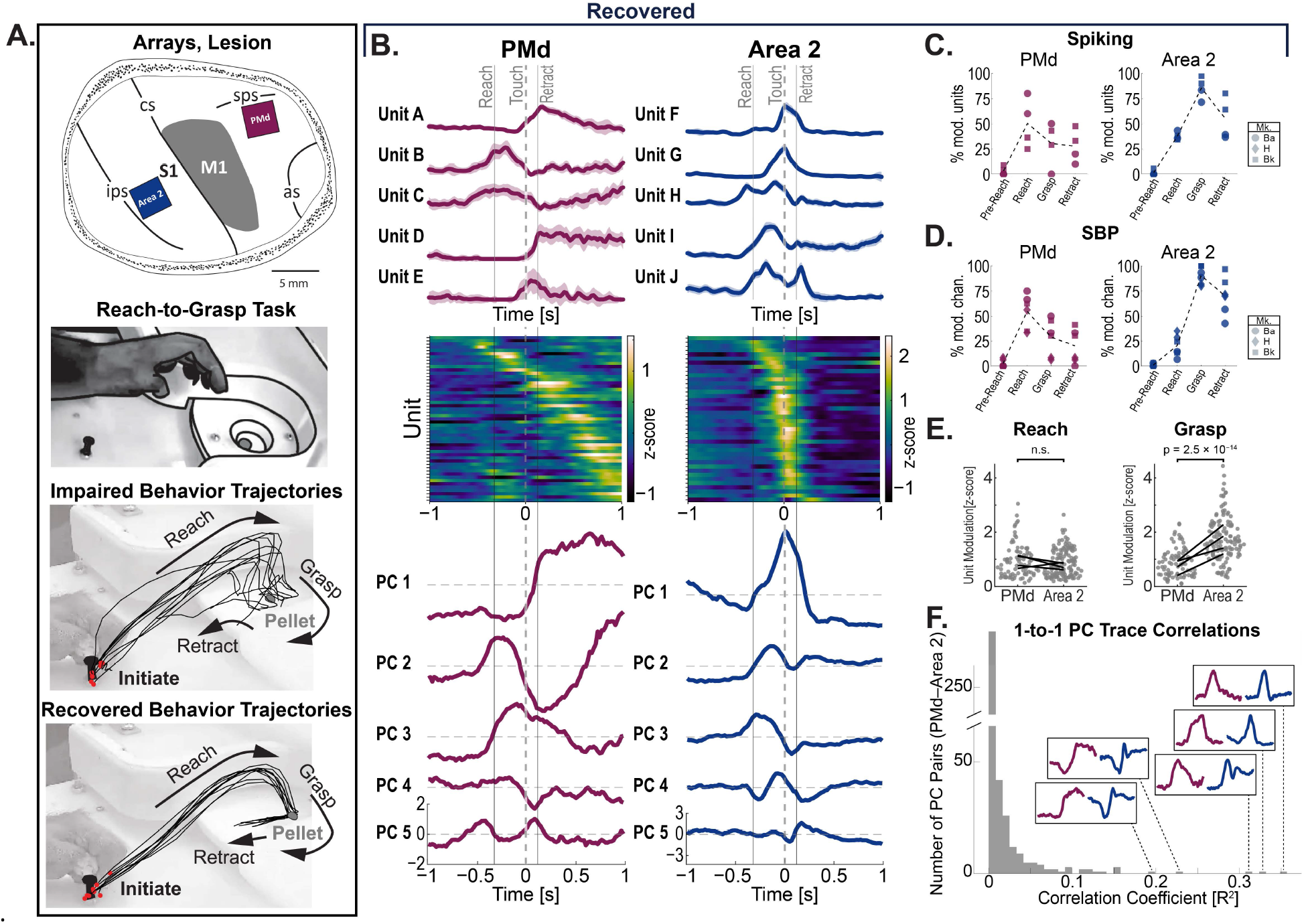
Activity during reach-to-grasp actions in PMd and Area 2. Local neural activity in PMd and Area 2 of parietal cortex each show reach-to-grasp modulation after stroke recovery. Single-area PMd and Area 2 responses during reach-to-grasp movement after stroke recovery. (A)Methods. Top: Implant locations and M1 lesion extent. Row 2: Reach-to-grasp task. Row 3: Impaired R2G trajectories post-lesion. Row 4: Recovered R2G trajectories later post-lesion. (B)Example extracellular electrophysiology unit activity from one session during R2G. Trial-averaged and aligned to pellet touch onset (time = 0). Data from PMd (left column) and Area 2 (right column). (Top) Trial-averaged spiking activity traces (example units). (Middle) Trial-averaged spiking activity (all units; one unit per row). (Bottom) Trial-averaged traces of PCs (PCs built on per-trial data). (C, D) Task-modulation of units (left) and spiking-band power (right). (C)Percentage of units modulated during different R2G epochs. (D)Percentage of SBP channels modulated by different R2G epochs. (E)Per-unit epoch modulation magnitudes comparing PMd to Area 2 spiking units (n = 2 animals; two sessions each). Per-session median modulations shown (lines). Statistics: Kolmogorov–Smirnov tests comparing epoch modulation of PMd versus Area 2 units for reach onset (“Reach”) and grasp onset (“Grasp”) epochs (task-modulated units only). (F)Pairwise correlation between PMd and Area 2 principal components with example PC traces. Distribution of Pearson correlation coefficients (R2) of trial-averaged PC traces calculated between PMd–Area 2 PC pairs (all combinations of top 10 PCs from each area). Trial-averaged PC traces for corresponding highly-correlated pairs are shown in insets (Left/maroon: PMd, Right/blue: Area 2).

During the same surgery, one 64-channel microwire electrode array was implanted in PMd and another in Area 2 (Figure 1A, Panel 1). The PMd array was anterior to the lesioned M1 area. Area 2 was chosen because it contains both cutaneous and proprioceptive information^11–16^ and is strongly connected with PMd through intermediaries including M1 and area 5 of posterior parietal cortex—in addition to sparse direct connections.^27^ Of the two animals with isolated spiking units on both the PMd and Area 2 arrays (Ba and Bk), after full recovery spiking activity showed substantial task-related modulation in line with previous findings.^28^ PMd units showed relatively heterogeneous firing patterns (Figure 1B, Left Column) with peak firing distributed throughout the duration of movement (Figure 1B, Row 2), while Area 2 activity appeared to be temporally biased to grasping and pellet contact (Figure 1B, Right Column). Principal component analysis resulted in top principal components (PCs) that seemed to qualitatively mirror spiking activity patterns (Figure 1B, Row 3). As expected^28^, units in both areas showed reach-to-grasp modulation, with PMd units appearing to be modulated during the reach epoch and Area 2 during the grasp epoch (Figure 1C). This task modulation was mirrored in the spiking-band power (SBP, 300-1000 Hz) activity (Figure 1D). SBP is a common proxy for spiking activity in brain-computer interfaces.^29,30^ Among task-modulated units, there was no significant difference in modulation magnitudes between PMd and Area 2 during reach; however, Area 2 did show higher modulation magnitudes than PMd during grasp (two-sample Kolmogorov–Smirnov test, test statistic = 0.55, p = 2.5 × 10^-14^). In summary, spiking activity in three fully recovered animals showed task-modulated activity in PMd and Area 2 during reach-to-grasp behavior.

Notably, a subset of neural activity patterns in PMd showed qualitative similarities to patterns of activity in Area 2 (Figure 1F, insets). To quantify this, we compared single-trial PC activity traces in PMd to those in Area 2 in the same session using Pearson correlation. While the majority of pairs do not show high levels of correlation, a small subset of PMd–Area 2 PC pairs show correlation coefficients distinctly higher than the typical distribution (Figure 1F). Together with the reach-to-grasp modulation patterns, these observations suggest that recovered upper-limb related neural activity in PMd and Area 2 is both comprised of components that may be similar across areas and components that may be unique to each region, motivating an explicit decomposition of cross-area and within-area population dynamics.

### Cross-area PMd–Area 2 factors during upper-limb movement in recovered animals

Similar to previous studies that measured population activity across two distinct areas,^24,31,32^ we used a dimensionality reduction technique to find latent factors that are explicitly shared across two areas^24^ (DLAG, Figure 2A). DLAG is a two-area extension of Gaussian-process factor analysis (GPFA)^33^ that identifies signals shared between two brain areas (cross-area factors, CFs). Additionally, DLAG estimates within-area factors, WFs—thus isolating activity patterns that are unique to either PMd or Area 2. As with GPFA, DLAG estimates a unique Gaussian kernel width that represents the timescale of activity fluctuations for that factor. Additionally, DLAG fits a temporal offset, “lag”, for each CF that best matches the temporal offset between when the corresponding signal occurs in one area versus the other. This allows for an estimation of the directionality for each of the cross-area signals.

**Figure 2.**
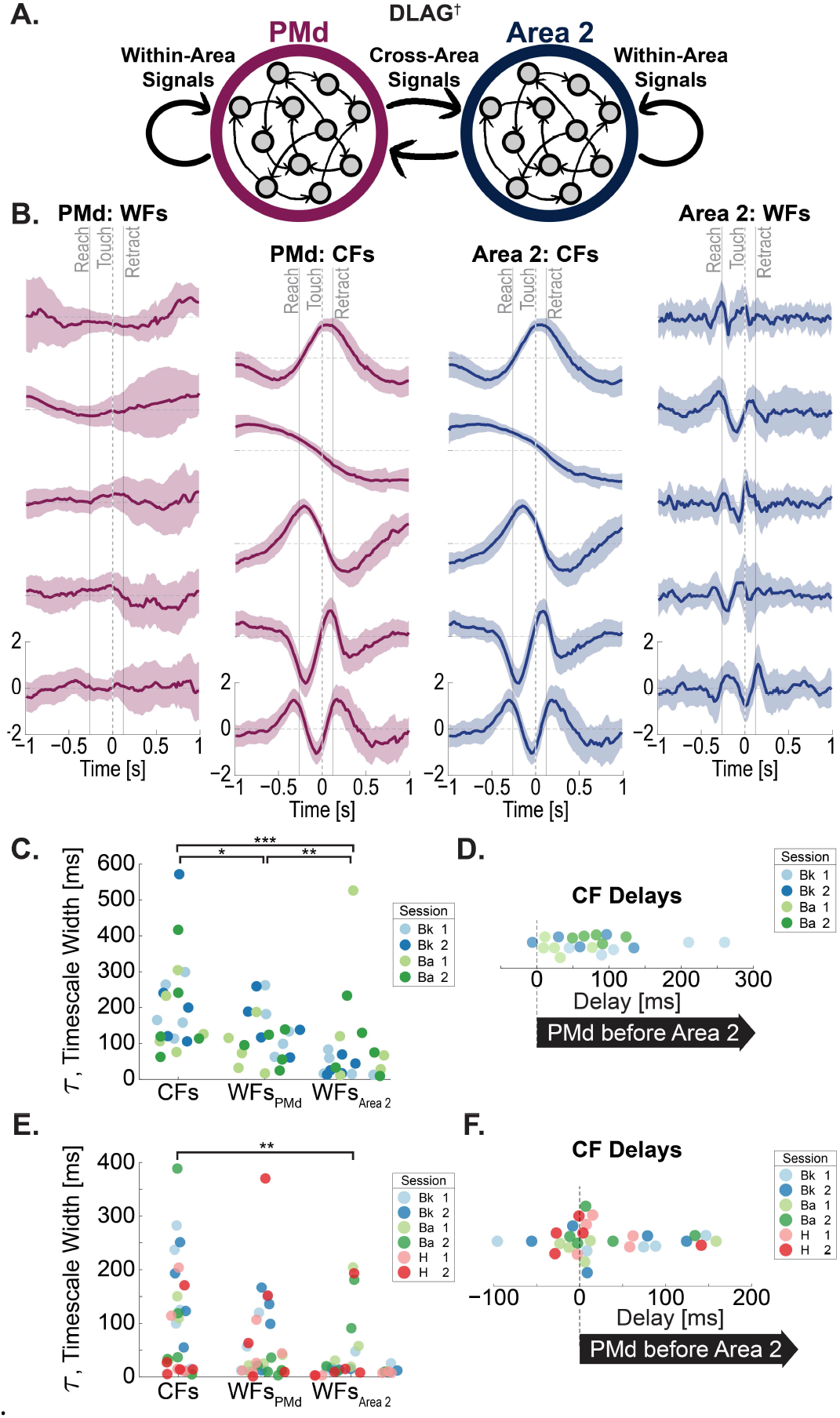
Cross-area PMd–Area 2 factors during reach-to-grasp behavior in recovered animals. (A)Illustration of dimensionality reduction technique DLAG†. Adapted from †Gokcen et al., 2022. (B)CFs and WFs from example session (Bk) using clustered spiking data. Median trace across trials; error ribbon: standard deviation. Column 1: WFs_PMd_; Column 2: CFs_PMd_; Column 3: CFs_Area 2_; Column 4: WFs_Area 2_. (C)Gaussian process timescale width (τ) for the CFs, WFs_PMd_, and WFs_Area 2_ for DLAG models fit on clustered spiking data in animals fully recovered from M1 lesion (n = 2 animals; two sessions each). Two-sided Wilcoxon rank sum tests shown differences in timescale widths; * CFs–WFs_PMd_: rank sum = 494, p = 0.0227; *** CFs–WFs_Area 2_: rank sum = 557, p = 2.5 × 10^-12^; ** WFs_PMd_–WFs_Area 2_: rank sum = 508, p = 0.0073 (D)The delay for cross-area factors. DLAG models fit on clustered spiking data in animals fully recovered from M1 lesion (n = 2 animals; two sessions each). Positive delay values indicate Area 2 activity followed corresponding PMd activity. (E)Gaussian process timescale width (τ) for the CFs, WFs_PMd_, and WFs_Area 2_ for DLAG models fit on low-SBP data in animals fully recovered from M1 lesion (n = 3 animals; two sessions each). Two-sided Wilcoxon rank sum tests shown differences in timescale widths; ** CFs–WFs_Area 2_: rank sum = 1114, p = 0.0029. (F)The delay for cross-area factors, low-SBP. DLAG models fit on low-SBP data in animals fully recovered from M1 lesion (n = 3 animals; two sessions each). Positive delay values indicate Area 2 activity followed corresponding PMd activity.

#### DLAG CFs and WFs

Using data from fully recovered animals, we fit DLAG models on single-session data of extracellular electrophysiology recordings from PMd and Area 2 recorded simultaneously during the reach-to-grasp task (Figure 2B). We input concatenated single-trial data of binned spiking rates, with each two-second trial epoch centered around the time of pellet touch into DLAG. Spiking data was spike sorted^34^ and included both isolated single-unit and multi-unit data. In fully recovered animals, DLAG found variance shared between the two simultaneously recorded arrays; as expected, these CF activations are mirrored between the two datasets but with a lag offset. Much like the spiking activity (Figure 1), these CF activity patterns show fluctuations throughout the R2G trial period that persist after trial-averaging, as seen in trial-averaged latent factor activations in an example session (Monkey Bk, Figure 2B, Columns 2 and 3). In contrast, PMd WFs showed activity fluctuations distributed broadly throughout the trial epoch (Figure 2B, Column 1). Area 2 WFs appeared to show sharper fluctuations that were concentrated during the reach and grasp periods (Figure 2, Column 4)—much like the temporal concentration of Area 2 spiking units (Figure 1).

PMd–Area 2 CFs showed significantly longer timescales than WFs from either region (medians, CFs: 162 ms; WFs_PMd_: 117 ms; WFs_Area 2_: 39 ms) (Figure 2C) (CFs–WFs_PMd_: rank sum = 494, p = 0.0227; CFs–WFs_Area 2_: rank sum = 557, p = 2.5 × 10^-12^)— meaning CFs embodied lower-frequency population activity patterns while Area 2 WFs had the most rapidly alternating patterns. Among WFs, Area 2 WFs tended to have shorter timescales than those in PMd (Figure 2C) (WFs_PMd_–WFs_Area 2_: rank sum = 508, p = 0.0073). This overall timescale pattern was mirrored in the models trained on low-SBP activity, albeit with CF timescale difference only reaching statistical significance comparing to WFs in Area 2 (Figure 2E) (CFs–WFs_Area 2_: rank sum = 1114, p = 0.0029). For the relative directionality of the CFs, the PMd component preceded the corresponding Area 2 component in most CFs fit to clustered spiking data (Figure 2D), with a less skewed distribution for models trained on low-SBP data (Figure 2F).

In summary, we found that a cross-area subspace can be detected between PMd and Area 2. The activity patterns of these cross-area factors showed a longer timescale than local WF factors; moreover, PMd was found to lead Area 2.

### Cross-area factors and kinematics

To better understand how these cross-area factors relate to behavior, we used kinematic tracking^35^ to investigate how CF activations correlate with R2G kinematics. Qualitatively, CF activity appeared to fluctuate in relationship to behavioral events (Figure 3A). In one trial (Figure 3A, Bottom), the animal fumbled its initial grasp attempt, which means it made two grasping motions over the pellet well (note two touch onsets, orange), and the arm therefore stayed extended for a longer duration, i.e. a longer ‘grasp’ epoch. During this extended grasp epoch, CF1 activity stays high during the grasp epoch, similar to the activity of CF1 during the grasp epoch of the previous trial. Similarly, CF5 activity shows a second, though smaller, positive bump with a time course matching the corresponding arm and hand motions made during the second grasp attempt. Moreover, consistent alignment to key kinematic timepoints (Figure 3A, vertical lines) appeared to persist throughout trials (Figure 3B).

**Figure 3.**
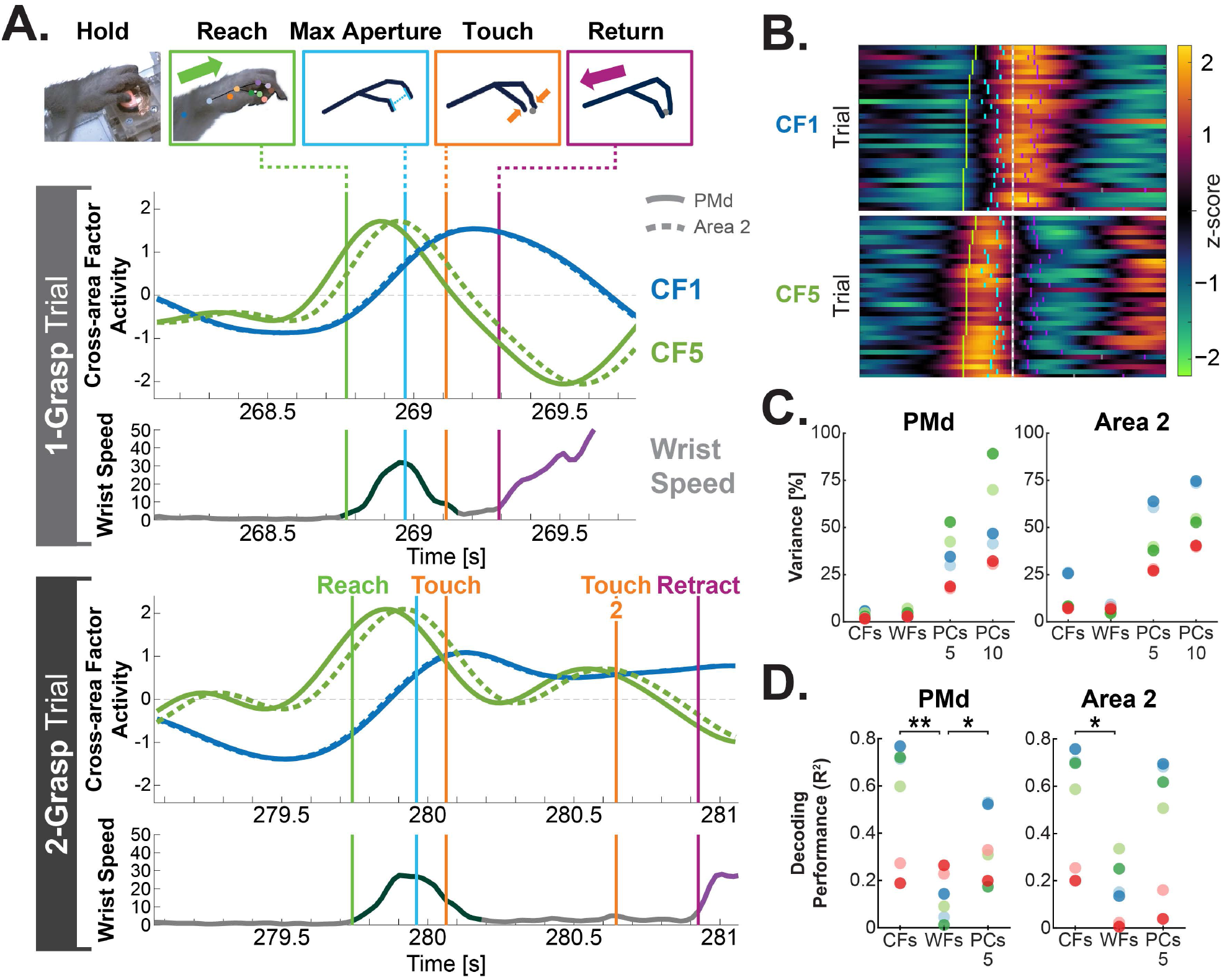
PMd–Area 2 factors contain substantial kinematic information despite accounting for a small portion of the total variance. (A)Single-trial latent factor activations (green and blue traces) for two CFs aligned to behavior. Both the PMd (solid) and Area 2 (dashed) version of each latent factor activation are shown. Key behavioral timepoints are denoted by vertical lines: reach start (green), point of maximum hand aperture (cyan), pellet touch (orange), and the instigation of retract at the end of the grasp (magenta). (Bottom) Time-aligned wrist speed (grey). Reach direction is annotated over the wrist speed to denote forward reach (dark green) and retract(purple). Example trials show (upper) a trial in which the animal successfully grasped the pellet on the first attempt and (lower) a trial in which the animal required two grasping attempts to successfully retrieve the pellet. (B)The same session as shown in A but showing all trials, with instantaneous CF latent value denoted by color scale. Trials are sorted by reach duration and aligned to the time of first pellet touch (time = 0). Key behavioral timepoints are denoted in single-trial tick marks: reach (green), maximum aperture (cyan), secondary grasp attempts (grey), and grasp finish (magenta). (C)Variance accounted for (VAF) by portions of DLAG models and PCA models fit on neural data in animals fully recovered from M1 lesion (n = 3 animals; two sessions each). DLAG models were fit on clustered spiking data for animals with clustered spiking units on both arrays (n = 2: Mk. Bk and Ba); low-SBP data was used on the animal that lacked adequate sortable units on the Area 2 array (n = 1: Mk. H). Each dot indicates the VAF by the DLAG CFs, DLAG WFs, top 5 PCs, and top 10 PCs for each session. (D)Decoding performance of reconstruction of wrist speed from neural latents. Cross-validated linear models fit on time-lagged duplicates of either PMd (left) or Area 2 (right) single-trial latents of CFs, WFs, or top 5 PCs. In PMd, both CFs and top 5 PCs had significantly higher decoding performance (coefficient of determination, R^2^) than WFs (rank sum: CF–WF, p = 0.0087 **; WF–PC_5_, p = 0.026 *, CF–PC_5_ = 0.18). In Area 2, CFs had significantly higher decoding performance than WFs (rank sum: CF–WF, p = 0.015 *, WF–PC_5_, p = 0.065; CF–PC_5_, p = 0.31).

To better quantify how these factors relate to kinematics, we used linear regression to quantify how well each area’s CF and WF activities could reconstruct wrist speed (Figure 3D) and hand aperture (henceforth “decoding”). As a comparison, we also calculated decoding performance using principal components (PCs) fit on single-trial data from either PMd or Area 2. Neural data consisted of single-trial clustered spiking data except for one animal that lacked adequate sortable units on the Area 2 array (Mk. H) in which case spiking-band power was used. For all cases, duplicate copies of temporally offset traces (lags: -180, -120, -60, 0 ms) were also included in the predictor variables to reduce the effect of temporal lags. Lag parameters were chosen through parameter search (Supplemental Figure 1) and were optimized for PC performance; they are in line with the published 100 to 160-ms range of optimal temporal offsets for motor cortex decoding.^36,37^ To provide parity with the Gaussian kernels that smooth DLAG CFs and WFs, a 40-ms width Gaussian smoothing kernel^33^ was applied to single-trial spiking data fed into PCA. This 40-ms kernel was also chosen by parameter search (Supplemental Figure 1). A non-parametric omnibus test confirmed there were significant differences between CFs, WFs, and top 5 PCs for both PMd and Area 2 (Kruskal–Wallis, PMd: p = 0.011, Area 2: p = 0.027) (Figure 3D). For hand aperture, decoding performance was lower across predictor types with decoding differences between CFs, WFs, and top 5 PCs more similar in value, so aperture decoding did not pass the omnibus test (Kruskal-Wallis, PMd: p = 0.39, Area 2: p = 0.062).

CFs were very predictive of wrist speed (Figure 3D) (R^2^_cross-validated_; PMd: 0.19-0.77, Area 2: 0.20-0.76), and strikingly PMd– Area 2 CFs were more predictive of wrist speed than all other predictor variables tested, though only the CF–WF comparisons reached significance (rank sum, PMd: CF–WF, p = 0.0087; WF–PC_5_, p = 0.026, CF–PC_5_, p = 0.18; rank sum, Area 2: CF–WF, p = 0.015, WF–PC_5_, p = 0.065; CF–PC_5_, p = 0.31). These results are surprising considering CFs and WFs accounted for a relatively small percentage of the total variance of the individual areas (Figure 3C) (PMd: CFs 2-6%, WFs 3-7%; Area 2: CFs 7-26%, WFs 5-9%)—especially compared to variance accounted for by top PCs (Figure 3C) (Top 5 PCs: PMd 30-53%, Area 2 38-64%).

### Cross-area factors emerge over recovery

Following a one-week post-surgery recovery period, two of the three animals demonstrated motor impairments (Ba and H). These two animals showed improved R2G task performance with time, recovering to within pre-stroke baseline levels after multiple weeks (Figure 4A). To examine changes in population-level activity over the course of recovery, we fit DLAG models on PMd–Area 2 data from individual R2G sessions collected in the weeks following recovery. We used the spiking-band power (SBP) per channel given that we sought to compare across sessions, and isolated spiking activity is known to emerge with recovery.^38^ Early after M1 stroke, the modulation of PMd–Area 2 CFs was weak (Figure 4B, left; see Figure 5 for single-trial examples). With recovery, modulation of PMd–Area 2 CFs increased (Figure 4B, right). The first two sessions after stroke with more than five rewarded trials were designated as “Early”, and the sessions in which R2G trial times were not significantly different than the pre-stroke behavior were designated as “Late”. We quantified the change in PMd–Area 2 CFs by a few physiologically relevant parameters detailed below.

**Figure 4.**
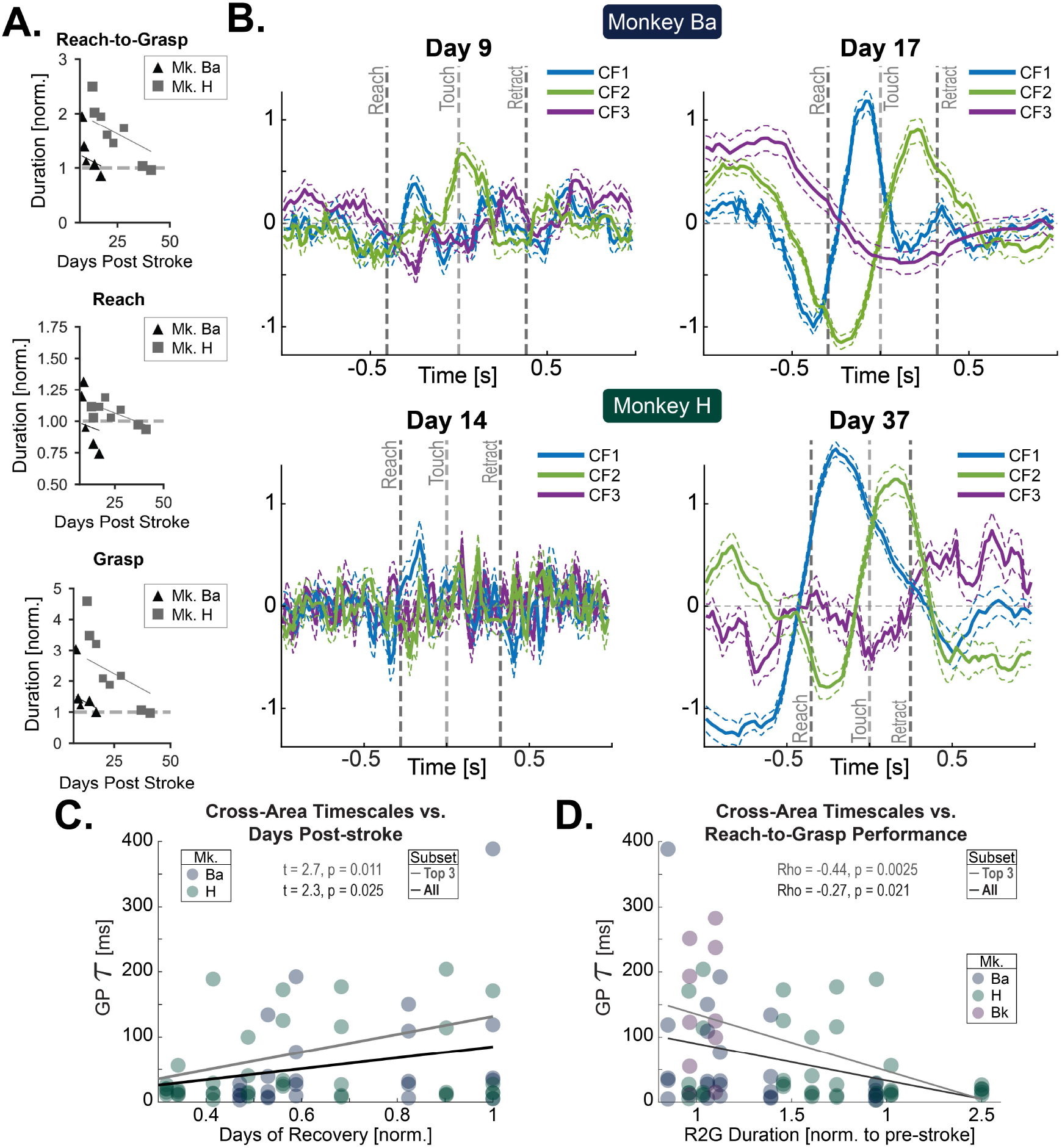
Longer timescale cross-area PMd–Area 2 factors emerge with behavioral recovery. (A)Behavioral improvements in two animals that showed post-lesion behavioral deficits. Baseline normalized R2G duration, normalized reach duration, and normalized grasp duration over recovery. Lines show LME fits with animal-specific intercepts. (B)Medians of similarly-timed trial CF traces (three of five CFs shown) for a session early after M1 lesion (left) and late into recovery (right) when the behavior had reached pre-lesion performance. Traces were “time warped” before median averaging to remove timing differences. Only successful trials were included. All DLAG results in Figure 4 are from models trained on low-SBP data. Both recovery animals shown (top and bottom) (n = 2). Error lines: standard error, SEM. CFs were ordered to group qualitatively similar latents. Trials aligned to pellet touch onset (time = 0). Single-trial trace examples are shown in Figure 5A. (C)CF Gaussian process widths (τ) per session over time (normalized days after lesion) for animals with recovery curves (n = 2). Lines show linear model fit for all CFs (black) and top 3 CFs (grey). (D)CF Gaussian process widths (τ) vs. normalized R2G duration for all animals (n = 3). Lines show linear model fit for all CFs (black) and top 3 CFs (grey).

**Figure 5.**
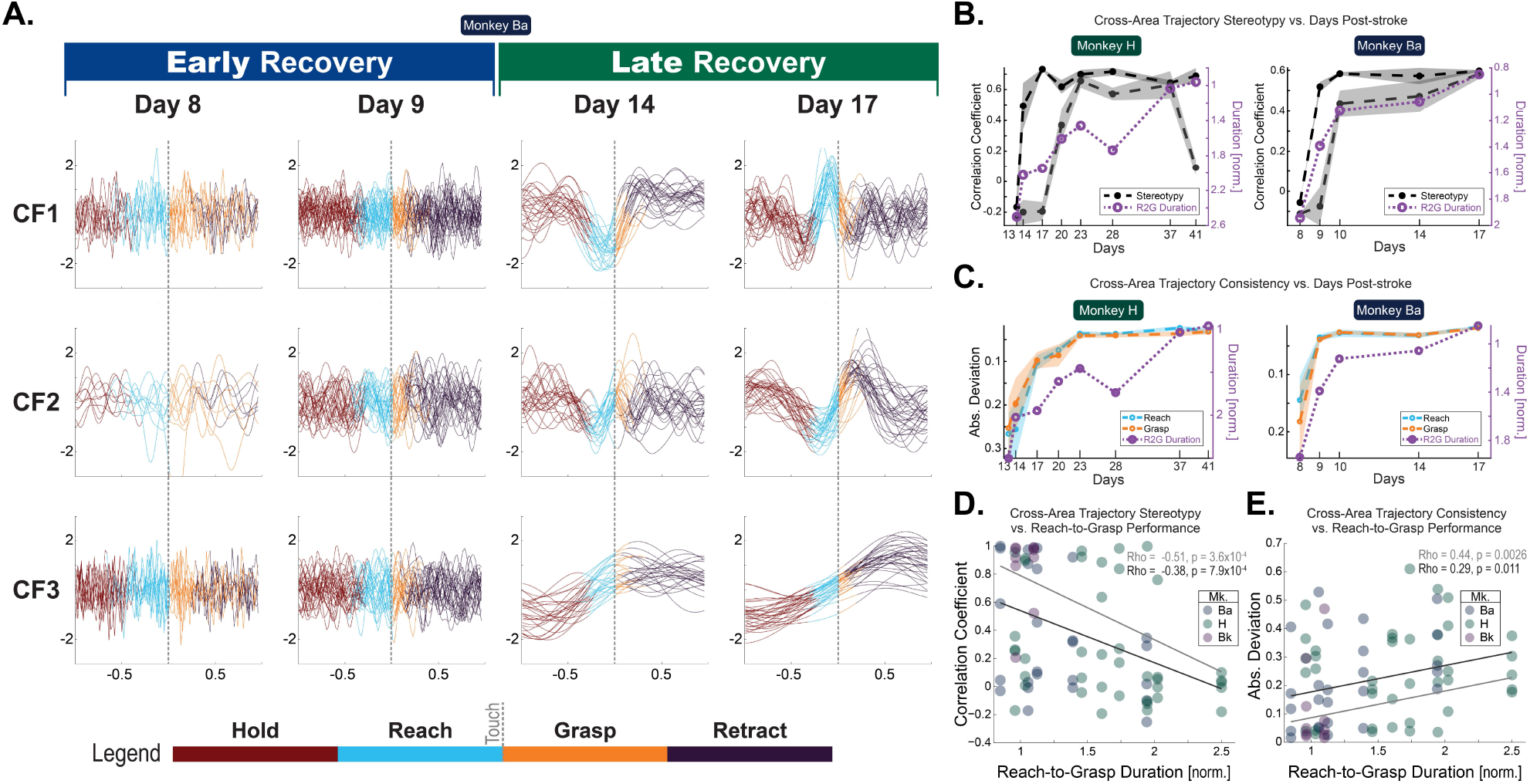
Trial-to-trial CF activation reliability improves with recovery. (A)Single-trial CF traces over recovery. All DLAG results in Figure 5 are from models trained on low-SBP data. Two sessions early after M1 lesion (left) and two late into recovery (right) when the animal’s behavior has reached pre-lesion performance (Mk: Ba). Single-trial traces of three of five CFs shown, with color indicating per-trial R2G epochs: pre-reach (maroon), reach (cyan), grasp (orange), and post-grasp retract(purple). Trials aligned to pellet touch onset (time = 0). CFs were ordered to group qualitatively similar latents. (B)CF trajectory stereotypy_position_ (black, grey) and R2G performance (purple) over time. Stereotypy_position_ is a R2G epoch-wide metric of trial-to-trial reliability of latent factor state, using the latent factor activation trajectory from reach onset to grasp finish. Per-trial stereotypy_position_ is calculated as the correlation of the single-trial trace vs. the mean trace of all other trials. The two most stereotyped CFs for each session are shown. Midlines: median per-trial stereotypy. Error: SEM. (C)CF trajectory consistency_velocity_ (cyan and orange) and R2G performance (purple) over time. Consistency_velocity_ is a timepoint-based metric of trial-to-trial reliability of latent factor state, using the instantaneous velocity at key R2G behavioral timepoints: Reach onset, Touch onset, and Grasp Finish onset (i.e. the color transitions in panel A). Per-trial consistency is calculated as the absolute deviation from mean velocity from all trials. Showing results for the most consistent CF for two R2G timepoint subsets: reach start (cyan) and grasp (orange); “grasp” is the mean of grasp start and grasp finish. Midlines: median per-trial consistency. Error: SEM. Normalized R2G duration per session (purple). (D)CF trajectory stereotypy_position_ vs. normalized R2G duration for all animals (n = 3). Template stereotypy is a R2G epoch-wide metric of trial-to-trial reliability of latent factor state, using the latent factor activation trajectory from reach onset to grasp finish. Per-trial template stereotypy is calculated as the correlation of the single-trial trace vs. the mean trace of all other trials. All CFs for each session are included. Lines show linear model fit for all CFs (black) and top 3 CFs (grey). (E)CF instantaneous consistency_velocity_ vs. normalized R2G duration for all animals (n = 3). All CFs for each session are included. Lines show linear model fit for all CFs (black) and top 3 CFs (grey).

#### Emergence of long-timescale CFs

Earlier we highlighted how, in fully recovered animals, PMd–Area 2 CFs are characterized by longer-timescale dynamics (Figure 2C, Figure 2E). In Early Recovery sessions, PMd–Area 2 CFs fit by DLAG had short timescales—reflecting the faster time-varying activations (Figure 4B, left). In contrast, PMd–Area 2 CFs in Late Recovery sessions, in which this animal’s performance did not differ significantly from pre-stroke performance, tended to have longer timescales—reflecting CF tendency for slower time-varying activations (Figure 4B, right). This trend of increased PMd–Area 2 CF timescales emerged over the course of recovery in the days and weeks following stroke (Figure 4C). Interestingly, tau significantly increased over time (LME with animal as random effect: t = 2.3, p = 0.025).

We also tested whether the emergence of long CF timescales correlated with the improvement of reach-to-grasp performance. We found that trial-averaged R2G performance correlated with longer PMd–Area 2 CF timescales (Figure 4D) (ρ_Spearman_ = -0.27, p = 0.021). Restricting to only those animals that showed a behavioral recovery curve (n = 2, Ba and H), we also found that PMd–Area 2 CF timescales significantly lengthened with improved trial-averaged R2G performance (LME with animal as random effect: t = -2.4, p = 0.020).

### Trial-to-trial consistency of cross-area PMd–Area 2 factor activity improves with recovery

Comparing behaviorally aligned single-trial CF activation traces for models built on Early versus Late, we found a marked qualitative improvement in CF consistency across trials (Figure 5A). To quantify this trial-to-trial consistency, we used two metrics: trial-wide stereotypy and timepoint consistency. Since CF activations in fully recovered animals were characterized by reliable traversals through similar positions in CF state space across trials, we used the reliability of the R2G epoch-wide latent factor activation trajectories for the first trial-to-trial reliability metric (henceforth “stereotypy_position_”). Specifically, a given trial’s stereotypy_position_ was calculated as the correlation of the single-trial trace versus the mean trace of all other trials.^19,21^ To reduce the possibility of performance speed variations influencing stereotypy_position_ results, only kinematically matched trials were included in its calculation, and those similarly timed trial traces were subsequently time warped to better align reach-to-touch and touch-to-retract epochs.

Since key kinematic moments appeared to align not only with the instantaneous state of the CF but also with the nature of the ongoing *change* of the CF state, we used the instantaneous *velocity* of each CF’s activity at key R2G timepoints for the second metric (henceforth “consistency_velocity_”)—velocity consisting of the first derivative of the CF activity trace. Specifically, a given trial’s consistency_velocity_ was calculated as the absolute difference of the velocity to the mean velocity across trials for key reach and grasp transitions: “reach”, “touch”, and “retract”. Since the use of instantaneous kinematic timepoints does not necessitate restriction to similarly timed trials, all trials with a successful pellet retrieval were included in consistency_velocity_ calculation.

Both stereotypy_position_ (Figure 5B) and consistency_velocity_ (Figure 5C) improved over days of recovery. Intriguingly, this improvement appeared to qualitatively coincide with R2G performance. Indeed, trial-averaged R2G performance per session significantly correlated with improved stereotypy_position_ (Figure 5D) (ρ_Spearman_ = -0.38, p = 7.9 × 10^-4^) and consistency_velocity_ (Figure 5E) (ρ_Spearman_ = 0.29, p = 0.011) when including data from all three animals. Restricting to only those animals that showed a behavioral recovery curve (n = 2), we found that PMd–Area 2 CF stereotypy_position_ significantly improved with better trial-averaged R2G performance (LME with animal as random effect: t = -3.0, p = 0.0044). CF consistency_velocity_ fell just short of significance (t = 1.9; p = 0.064) when including all 5 CFs. Restricted to just the most consistent 3 of 5 CFs, the R2G performance–consistency_velocity_ was significant .

In summary, the trial-to-trial reliability of PMd–Area 2 cross-area factor activity improved over the days and weeks following the loss of the shared cortical neighbor, M1. These improvements in trial-to-trial reliability were correlated with reach-to-grasp function.

### Within-area changes did not account for behavioral recovery

One of the strengths of DLAG is that it simultaneously fits CFs and WFs. In the previous sections, we have shown how slower timescale, consistent CFs emerge over the course of recovery from M1 stroke. We also investigated whether WFs were also correlated with recovery.

There was no significant change in PMd or Area 2 WF timescales (tau) over the course of recovery in the days and weeks following stroke (LME with animal as random effect; PMd: t = 1.2, p = 0.25; Area 2: t = 1.0, p = 0.32). Similarly, there was no significant change in PMd or Area 2 WF tau values relative to per-session R2G performance (LME with animal as random effect; PMd: t = -0.79453, p = 0.43; Area 2: t = -1.4, p = 0.18) (Supplemental Figure 2A).

#### Within-area: Stereotypy & consistency over time

Including data from all three animals, trial-averaged R2G performance per session had no correlation with WF stereotypy_position_ (PMd: ρ_Spearman_ = -0.16, p = 0.17; Area 2: ρ_Spearman_ = -0.18, p = 0.11) (Supplemental Figure 2C) or consistency_velocity_ (PMd: ρ_Spearman_ = 0.15, p = 0.21; Area 2: ρ_Spearman_ = -0.09, p = 0.44) (Supplemental Figure 2D) for neither PMd nor Area 2. Restricting to only those animals that showed a behavioral recovery curve (n = 2), again there was no correlation with WF stereotypy_position_ (LME with animal as random effect; PMd: t = -1.1, p = 0.26; Area 2: t = -1.3, p = 0.19). However, while there was no trend with PMd WF consistency_velocity_ (LME with animal as random effect; PMd: t = 1.3, p = 0.19), there did appear a significant trend for Area 2 WF consistency_velocity_ versus trial-averaged R2G performance per session (LME with animal as random effect; t = -2.4, p = 0.018). But importantly, this Area 2 WF consistency_velocity_ trend was inverted (slope = -0.12) compared to the significant trend found for CFs (slope = 0.07), i.e. Area 2 WF instantaneous velocity at key behavioral moments was *less* consistent in sessions with better behavioral performance.

In summary, WF timescales neither lengthened nor shortened over the weeks of recovery. In terms of trajectory reliability, there were almost no clear trends seen regarding stereotypy_position_ or consistency_velocity_ relative to changes in R2G performance.

## Discussion

We provide the first population-level analysis of cross-area motor–somatosensory dynamics during changes in reach to grasp function over recovery. Early after M1 hand/arm injury, cross-area activity between perilesional PMd and Area 2 was weak and inconsistent. Over weeks of recovery, cross-area activity became of longer timescale and was increasingly stereotyped across trials. These findings suggest that the re-emergence of a shared motor–sensory subspace is important for recovery of prehension after stroke.

### Slow timescales and cross-area coordination

Smooth neural trajectories with low frequency rotational dynamics have been proposed to be key for execution of motor behaviors.^39,40^ Such dynamics are thought to support the generation of quasi-oscillatory temporal patterns that unfold reliably over time,^40^ likely rendering motor output more robust to noise and reducing trial-to-trial variability.^41,42^ These quasi-oscillatory patterns may also facilitate effective communication with downstream motor structures, including the striatum and the musculature, by providing temporally organized drive signals.^43–46^ Such patterns appear to be ‘condition-invariant’ and may be particularly important for movement timing.^47^ Slow timescale dynamics are perhaps a natural consequence of recurrent network architecture, where recurrent connectivity stabilizes low-dimensional population trajectories.^40^ Importantly, this recurrence may involve subcortical structures such as the thalamus.^48,49^ The thalamus possesses recurrent connections with both motor^50,51^ and somatosensory areas^52–54^, with evidence that these distributed loops further shape the temporal structure^55^ and the slow timescale of motor-related population activity.

Consistent with these ideas, we observed that cross-area PMd–Area 2 signals exhibited quasi-oscillatory temporal structure as animals recovered reach-to-grasp function (Figure 4). Such smooth, low-frequency population dynamics may provide a noise-robust communication channel, ensuring that behaviorally relevant information is reliably transmitted across sensorimotor circuits. One candidate for such information is corollary discharge, where accurate representation of motor intent or state is essential for appropriate interpretation of incoming sensory feedback.^56^ Mismatches between motor-related signals and sensory input would be expected to degrade sensorimotor integration and impair skilled performance. In line with this interpretation, reliable and stereotyped cross-area dynamics emerged only as skilled behavior recovered (Figure 5). Future work could investigate these possibilities further by building upon previous insights derived through computational approaches,^40,42,57^ perhaps by studying cross-area signals between sparsely connected recurrent neural networks.^57^

### Premotor–somatosensory coordination may enable recovery of dexterity

Coordinated motor–sensory signals are likely to be most important for prehension and skilled object interactions, such as picking up a small pellet as in our case. One possibility is that such precise object interactions require continuous alignment between internally generated motor commands and sensory feedback about limb and object state. Rather than operating as isolated encoders, motor cortical populations may function as generators of movement^41,42,58–61^ whose outputs must be dynamically reconciled with somatosensory information to support precise prehension. In dexterous behaviors, such as reach-to-grasp and object manipulation, accurate timing and scaling of muscle activation likely depend on reliable communication of motor intent, predicted sensory consequences, and ongoing sensory feedback.

Coordinated population dynamics across these regions may therefore provide a low-noise^41,42^, temporally structured substrate for integrating tactile and proprioceptive signals, enabling rapid adjustments to grip force, finger positioning, and object stability. Disruption of this coordination would be expected to impair fine motor control even when gross movement remains intact, whereas restoration of structured motor–sensory dynamics would support the recovery of dexterity. Thus, coordinated motor–sensory population signals may represent a fundamental mechanism by which the nervous system achieves robust, flexible control of object-directed actions.

### Cross-area activity and variance accounted for

One important consideration is the variance accounted for by cross-area factors relative to principal components (Figure 3C). In light of growing recognition that a large fraction of population activity reflects condition-independent processing,^47^ it is plausible that, in recurrent cortical networks, much of the observed variance arises from internally generated dynamics^40,60,62^ that are not directly task relevant.^47^ Under this framework, task- or movement-related signals may occupy only a small subspace of overall neural activity.^47,63^ Signals within this cross-area subspace may therefore exert a disproportionate influence on behavior despite explaining relatively little variance. Consistent with this view, the cross-area factors identified here were predictive of kinematic state and recovery (Figure 3), suggesting that they capture a compact, behaviorally important communication channel rather than more dominant local dynamics. Thus, low variance explained may reflect selective routing of information across areas rather than weakness of the signal, further highlighting the importance of isolating functionally relevant population dimensions beyond total variance metrics.

### Implications for neuromodulation

A key goal of understanding the neurophysiology of recovery is to develop tailored brain stimulation to improve recovery.^45^ Our findings suggest that recovery after stroke is supported by the re-emergence of coordinated population dynamics operating at slow timescales, as reflected in smoother, low-dimensional cross area activations. Interestingly, rotational dynamics in the intact M1 also appear to evolve at frequencies in the delta range^39^ (∼1–4 Hz) and are closely aligned with similar LFP fluctuations.^46^ This potential link between slow-timescale population rotations and LFP activity has also motivated neuromodulation strategies that target low-frequency dynamics (∼3 Hz) in order to improve neural co-firing and motor function.^20,21,23^ Our results suggest that such stimulation may also serve to stabilize inter-areal communication. What might be the circuit basis for slow timescale modulation? One possibility is that thalamic inputs to cortical regions are important for setting the timing of coordinated fluctuations.^44,64^ Together, these results argue that effective neuromodulation strategies could target distributed slower timescale population dynamics.

### Implications for sensorimotor integration

PMd–Area 2 cross-area factors suggested a skew towards signals occurring first in PMd and subsequently in Area 2 (Figure 2D, F). This bias is potentially surprising given the theories surrounding the role of sensorimotor feedback on motor control, including but not limited to optimal feedback theory, and previous evidence showing that motor cortex can be modulated by sensory expectation.^2^ We highlight two possible reasons for this skew. Firstly, this heavily PMd-then-Area 2 skew could be a result of our behavioral task and animals having to compensate for injury. Previous work has suggested that response order and latencies among cortical nodes of the sensorimotor network can change depending on the context of the task or feedback.^65,66^ We would expect alternative tasks that specifically manipulate feedback and generate controlled error conditions, such as Kinarm-based perturbation tasks,^2,66^ could be better suited to tease apart directionality. Secondly, we recorded from only two regions, PMd and Area 2, while many additional subcortical^67,68^ and cortical^27,69,70^ areas also contribute to feedback control^66,71^ and sensorimotor integration.^69,72^ Future work could record simultaneously from additional sensorimotor nodes and leverage novel tools, such as multi-area variants of DLAG,^73,74^ to better disentangle directionality in a complex high-interconnected system.

### Limitations

This study does not explicitly dissociate passive sensory input from active sensation, as we did not independently track tactile afferent signals during reach-to-grasp behavior. Instead, our analyses focus on coordination between motor and somatosensory populations during active movement and pellet contact, in which sensory signals are likely inherently shaped by movement. It therefore remains possible that altered sensory feedback contributes to the observed changes in cross-area coordination following stroke. However, we did not lesion somatosensory cortex, and local activity within Area 2 was not predictive of behavioral recovery, as would be expected if loss of sensory representation were the primary driver of impairment. Moreover, our analysis was only based on correct trials that were kinematically matched, where the pellet was successfully gripped. However, future studies that explicitly track traditional somatosensory signals, including tactile afferent encoding during controlled passive sensory stimulation, will be important for further dissociating changes in primary sensory representation from alterations in active sensorimotor coordination.

## Acknowledgements

We thank Evren Gokcen and Nikhilesh Natraj for helpful discussions and suggestions regarding analytical methods, and M.J. Lemoy and K. Stotts for expert surgical assistance. This research was funded by the NIH (NS112424 and NS117406 to K.G.) and was additionally supported by the Weill Institute for Neuroscience at UCSF and the Weill Neurohub Pillars Program. The Weill Neurohub Fellows Program also supported this project (to I.S.H).

## Methods

### Macaques

Experiments were conducted in compliance with the NIH Guide for the Care and Use of Laboratory Animals and were approved by the University of California, Davis Institutional Animal Care and Use Committee. Three (two male and 1 female) adult rhesus macaques (*Macaca mulatta*) were used in this study. Animals were 5–6 years old and 9.6–14.3 kg. Two animals were pair-housed while one was singly housed at the time of the data presented here. Animals were housed in rooms with lights that turned on at 6 am and off at 6 pm.

### Reach-to-Grasp Task

Macaques were trained to perform a reach-to-grasp (R2G) pellet retrieval task.^21,75^ To initiate a trial, animals touched a touch-sensitive peg. After holding the peg for a random hold time (uniformly distributed between 0.5–0.8 s), an automated apparatus released an edible pellet from an automated pellet dispenser (Lafayette Instrument Company, Lafayette, IN) into one of five “wells”. Wells were 5.9 mm deep with diameters of either 13 mm, 19 mm, 25 mm, 31 mm, or 37 mm. The smaller wells served to make pellet retrieval more challenging for the animals. Animals had 5 s to retrieve the pellet from the well before the apparatus would rotate the well out of view. Animals performed 10–100 trials per day, depending on level of impairment.

Two cameras (CM3-U3-13Y3C-CS, Teledyne FLIR, Richmond, BC, Canada) recorded a side and top view of the animal performing the task at 50 frames per second. Camera frames and touch sensor activity were synchronized using the electrophysiology recording system. Offline, these videos were used both for manual behavioral timepoint annotation and markerless kinematic tracking (see “Kinematic Tracking“).

#### Behavior Timepoints

Key behavioral timepoints for each R2G trial were identified to enable epoch segmentation, calculation of performance metrics, and visual references in figures. Reach onset, touch, and retract were determined via manual annotation by reviewing video frames. Maximum aperture was derived from coordinates determined by markerless kinematic tracking (see “Kinematic Tracking“). “Reach onset” was the video frame at which the animal first initiated forward motion from the hold point. “Touch” was the video frame at which the animal first made contact with the pellet; this co-occurred with the hand arriving at the well. “Retract” was the point at which the pellet was in control and the animal had begun retracting its arm; this is approximately equal to the completion of the grasp, and the hand is still above the well at this time. “Maximum aperture” was timepoint between “reach onset” and “touch” at which the distance between the thumb and index finger (in pixels) was maximal (see “Kinematics” for hand aperture estimation details).

### Surgery

#### Implantation

After reaching stable performance for the reach-to-grasp task, electrode implantation surgery was performed. Preoperatively, animals were sedated with ketamine hydrocholoride (10 mg/kg), administered atropine sulfate (0.05 mg/kg), prepared, and intubated. They were then placed on a mechanical ventilator and maintained on isoflurane inhalation (1.2%–1.5%). Animals were positioned in a stereotactic frame (David Kopf Instruments, Tujunga, CA) and administered mannitol (1.5 g/kg) intravenously prior to the craniotomy. An incision, craniotomy, and durotomy were performed over the lateral frontoparietal convexity of the right hemisphere, exposing the caudal region of the frontal lobe and the rostral region of the parietal lobe.

Each macaque was surgically implanted with two chronic microwire arrays: one in the forelimb area of dorsal premotor cortex and one in Brodmann Area 2 within somatosensory cortex.^21^ Area 2 was chosen because it contains both cutaneous and proprioceptive information.^11–13^ Arrays were 64-channel tungsten microwire multielectrode arrays (Tucker-Davis Technologies, Alachua, FL) with an 8-by-8 channel arrangement and 500-by-375 µm orthogonal channel spacing. Arrays were inserted 2 mm into the cortex using a micromanipulator mounted to the stereotax. Array ground and reference wires were tied to titanium skull screws (Gray Matter Research, Bozeman, MT) that were in contact with the dura, with dural contact confirmed during surgery and post-mortem. Following surgery, animals were administered analgesics and antibiotics and were carefully monitored post-operatively for 7 days.

#### Lesion

During the same surgery as array implantation, animals underwent surgical induction of stroke localized to the forelimb area of primary motor cortex, specifically the caudal bank of the central sulcus, by surface vessel occlusion followed by subpial aspiration.^21,26^ The lesion perimeter was set using anatomical landmarks, extending dorsally to a horizontal level that included the precentral dimple (the lateral-most portion of the M1 leg area) and ventrally to the central sulcus genu (the dorsal-most portion of the M1 face area). After aspiration, a dural flap was sutured to cover the lesioned area. Six to eight 3-mm or 5-mm titanium screws inserted into the skull acted as structural support anchors for the array and a dental acrylic headcap, with the headcap serving both as additional support for the arrays and to seal the craniotomy. Each animal also had four to six additional skull screws with corresponding epicranial wire leads that served as stimulation screws^21^, but these data were not analyzed in this study.

### Electrophysiology

Raw neural signals were acquired at 24414.06 Hz using an amplifier (PZ5), signal processor (RZ2), and data storage system (RS4) (Tucker-Davis Technologies, Alachua, FL). Offline, two types of neural signals were derived from these recordings: neural spiking unit events and low spiking-band power (low-SBP). Low-SBP is used in the brain–computer interface field since it strongly covaries with spiking activity.^30^ We leveraged low-SBP in the recovery portion of this analysis because 1) it enables us to make comparisons across days without the confound of spiking unit identities changing between days,^76^ and 2) one of two animals with a prolonged behavioral recovery curve (Monkey H) lacked substantial sortable spiking units on its Area 2 array but had low-SBP activity.

#### Neural Activity Calculation

To sort spiking units, raw neural signals were first median subtracted (median computed on each bank of 16 channels in the array) to reduce motion artifacts and external noise. Spiking units were then isolated from these median-subtracted signals using Kilosort3^34^ and curated in Phy^77^ to remove noise units. All neural units above the minimum firing rate threshold were used for analysis, meaning both single-unit and multi-unit activity were included. The minimum threshold was set to a session-wide firing rate of 1 Hz—the one exception being the last session for one animal (session “Ba B”) for which a lower 0.5-Hz threshold was used to due to signal degradation of one of the arrays. Per-unit spike counts were binned into 20-ms bins and then normalized by the average firing rate of the corresponding unit across the entire recording session.

Low spiking-band power bins, “low-SBP”, were calculated from local field potential traces acquired at 1,017.25 Hz. Offline, these voltage traces were high-pass filtered at 240 Hz using a 5^th^-order Butterworth filter. These voltages were then median subtracted (median computed on each bank of 16 channels in the array) to reduce motion artifacts and external noise. Voltage traces were then z-scored. To calculate low-SBP per channel, we used a multi-taper method^78^ using Chronux (function: *mtspecgramc*) with 20-ms bin width and 20-ms sliding window width. Power values were averaged across the frequencies of the resulting estimated spectrum that fell within our low-SBP range: 300–500 Hz.

#### Neural Activity Postprocessing

For both spiking units and low-SBP, analysis was run on per-trial data—specifically on two-second epochs surrounding each trial, with the center of each two-second epoch aligned to the moment the pellet was first touched. Per-unit (or per-channel, for low-SBP) neural activity values were subsequently square-root transformed,^33^ demeaned by trial average per unit,^24^ and z-scored per unit. Trials in which the animal did not attempt the task were excluded from analysis.

Note that Gaussian smoothing was applied to neural activity in some but not all cases. Specifically, no smoothing was applied before feeding neural values into DLAG, in line with the method’s requirements.^24^ However, a 40-ms Gaussian smoothing kernel^33^ was applied to data fed into PCA or not passed to any dimensionality reduction method, with this smoothing step occurring between the aforementioned square-root transform and demean-by-trial steps.

### Regression and Linear Mixed Effects

We used linear mixed-effect (LME) models where possible, with animal as a random effect. One of three animals (Monkey Bk) had no recovery curve, i.e. R2G performance was already at baseline levels by our first neural recordings. This precludes the use of LME with normalized behavioral recovery as a predictor. In cases where all three animals were included, Spearman correlation was used instead of LME. LME models were implemented in MATLAB version R2021a using the “fitlme” function from the Statistics and Machine Learning Toolbox (MathWorks, Natick, MA).

### Consistency and Stereotypy

We used two metrics to quantify trial-to-trial reliability of time-varying DLAG factors: trial-wide stereotypy, “stereotypy_position_”, and timepoint consistency, “consistency_velocity_”.

Stereotypy_position_ of a given trial was the Pearson’s correlation coefficient calculated as the single-trial trace correlation to a “template” consisting of the mean trace of all other trials, and the median correlation for each session was reported.^19^ Only trials in which the animal successfully grasped the pellet on the first grasp attempt were included in this calculation. To further reduce the effect of different performance speeds on stereotypy_position_ calculation, only similarly timed trials were included in calculation, as well. Specifically, a trial had to be within ±0.1 s of both the median reach-to-touch duration (“reach onset” to “touch” timepoints) and the median touch-to-retract duration (“touch” to “retract” timepoints) for the session.

Consistency_velocity_ uses the first derivative of a single-trial neural activity trace at trial-specific behavioral timepoints. The addition of a first-derivative metric (i.e. “velocity” if zeroth derivative is “position”) highlights the ongoing change of neural state relative to behavior. Values were collected for three behavioral timepoints: reach onset, touch, and retract (see “Behavior Timepoints“), where the value for a trial is the absolute difference of the instantaneous velocity to the mean instantaneous velocity across all trials in that session, and the median value for each session was reported. The first derivative was estimated as the difference between adjacent elements. Since some factors were consistent at one but not all timepoints, we used the “best” result among the timepoints for each factor (i.e. the lowest median absolute deviation from the mean) for analysis. All trials were included in consistency_velocity_ calculation since the metric is trial duration agnostic.

### DLAG

DLAG^24^ was used to find patterns of neural activity shared between PMd and Area 2 (cross-area factors, CFs) and isolate additional latent variables of activity unique to either PMd or Area 2 (within-area factors, WFs). Preprocessed single-trial neural activity bins (see Postprocessing (Spiking) and Postprocessing (Low-SBP)) from PMd and Area 2 were supplied to DLAG. In line with DLAG usage, neural activity was not smoothed prior to fitting of DLAG models, since fitting of Gaussian process widths is part of the model fitting process. To enable comparisons across days, DLAG models were requested to fit a set number of factors: five factors for CFs, WFss_PMd_, and WFs_Area 2_ each. The maximum number of expectation–maximization algorithm iterations allowed per DLAG model training was set to 10^6^. All other DLAG parameters were set to defaults.^24^

### Kinematic Tracking

Forelimb kinematics were derived offline from video recordings via markerless kinematic tracking using DeepLabCut,^35^ which returned pixel coordinates of tracked anatomical locations on the wrist and hand. Kinematic trajectories for Figure1A were built from these tracked coordinates.

Wrist speed was derived as the two-dimensional distance traveled between video frames. Wrist speed was calculated on three redundant tracked points on the wrist; the mean of these three speed traces was used for analysis. This was done to increase robustness to momentary tracking loss, such as due to occlusion or motion blur. Wrist speed was smoothed with a 5-sample Gaussian kernel.

Hand aperture was derived as the distance between the thumb and index finder. Thumb and index finger location was calculated using the mean of two redundant points on the palmar surface of each of the two digits. Again, this was done to increase robustness to momentary tracking loss due to occlusion or motion blur.

### Kinematic Decoding

The ability of different forms of dimensionality-reduced neural data to predict subsequent kinematic output was estimated by fitting linear regression models. Model predictors consisted of time-lagged duplicates of single-trial latents of the CFs, WFs, or PCs from either PMd or Area 2 of somatosensory cortex. Specifically, time-lagged duplicates consisted of four total copies: one real and three lagged in 60-ms steps into the past (maximum offset: 180 ms before behavior). PCs were built on single-trial bins smoothed with a 40-ms Gaussian kernel. These parameters were chosen via a parameter search (Supplemental Figure 1) to optimize PC decoding performance, while also balancing for CF and WF performance. For decoding, lagged duplicates were included to avoid the confound of exact alignment affecting the results. This is especially important given that DLAG CFs inherently have lags. Wrist speed and hand aperture traces (see “Kinematic Tracking“) were matched to neural bin timeframe using linear interpolation, and model fitting was performed using the MATLAB function “fitlm”. Timepoints with missing values were excluded from decoding analysis. Results were cross-validated using repeated hold-out validation with an 80/20 split and 30 iterations.

For comparing in cases of fully recovered animals (Figure 3), DLAG (CFs and WFs) and PC models were built using clustered unit spiking activity for animals that had adequate spiking data (n = 2, Monkeys Ba and Bk), while the spiking-band power was used for the other animal (n = 1, Monkey H). Stats on decoding differences were run on decoding performance for three 5-dimension data types: CFs, WFs, and the top 5 PCs (“PC5”). Two late-recovery sessions for each of the three animals were included in the statistics, with separate tests being run on PMd and Area 2 subsets. Omnibus tests, Kruskal–Wallis tests, were run first to test for any difference between CF, WF, and PC_5_ decoding performance for wrist speed and hand aperture. If the omnibus null hypothesis was rejected (p < 0.05), post-hoc tests (two-sided Wilcoxon rank sum tests) were run for each individual comparison.

### Spiking Unit and SBP Channel Modulation

Per-unit and per-channel task and epoch modulation (Figure 1) was determined using the trial-averaged activity trace trace. After processing the individual spiking or power bins (*Neural Activity Postprocessing*), similarly-timed trials were “time warped” (*Time Warping of Activity Traces*) and trial averaged. Trial-averaged traces that deviated outside +/-1 standard deviation anywhere within a two-second window around pellet touch were considered “task modulated”. Units or channels were “epoch modulated” if they deviated outside +/-1 standard deviation during the corresponding 200-ms epoch. “Pre-reach” epochs 500 to 300 ms before reach onset. “Reach” was 100 ms before to 100 ms after reach onset. “Grasp” was 50 ms before to 150 ms after pellet touch, i.e. “touch”. “Retract” was 0 to 200 ms after retract onset, i.e. “retract” (see *Behavior Timepoints*).

The per-unit or per-channel modulation value was the absolute value of the maximum deviation of the trial-averaged trace during the epoch in question. These values were pooled across sessions and subjects to gauge whether PMd and Area 2 activity units or channels had different modulation patterns for key epochs with a two-sample Kolmogorov– Smirnov test (Figure 1).

### Time Warping of Activity Traces

Single-trial activity traces were temporally aligned, “time warped”, before trial averaging for certain illustrations and for spiking unit and SBP channel modulation calculations (Figure 1, *Spiking Unit and SBP Channel Modulation*). For time warping, only similarly timed trials were used. Specifically, trials had to be within 200 ms of both the median reach-to-touch and median touch-to-retract time durations for that session. Time warping was applied independently to before-touch and after-touch periods by linearly interpolating to align each trial to the median reach-to-touch and median touch-to-retract durations, respectively.

## Supplemental Figures

### SFig 1: Decoding parameter search

**Supplemental Figure 1.**
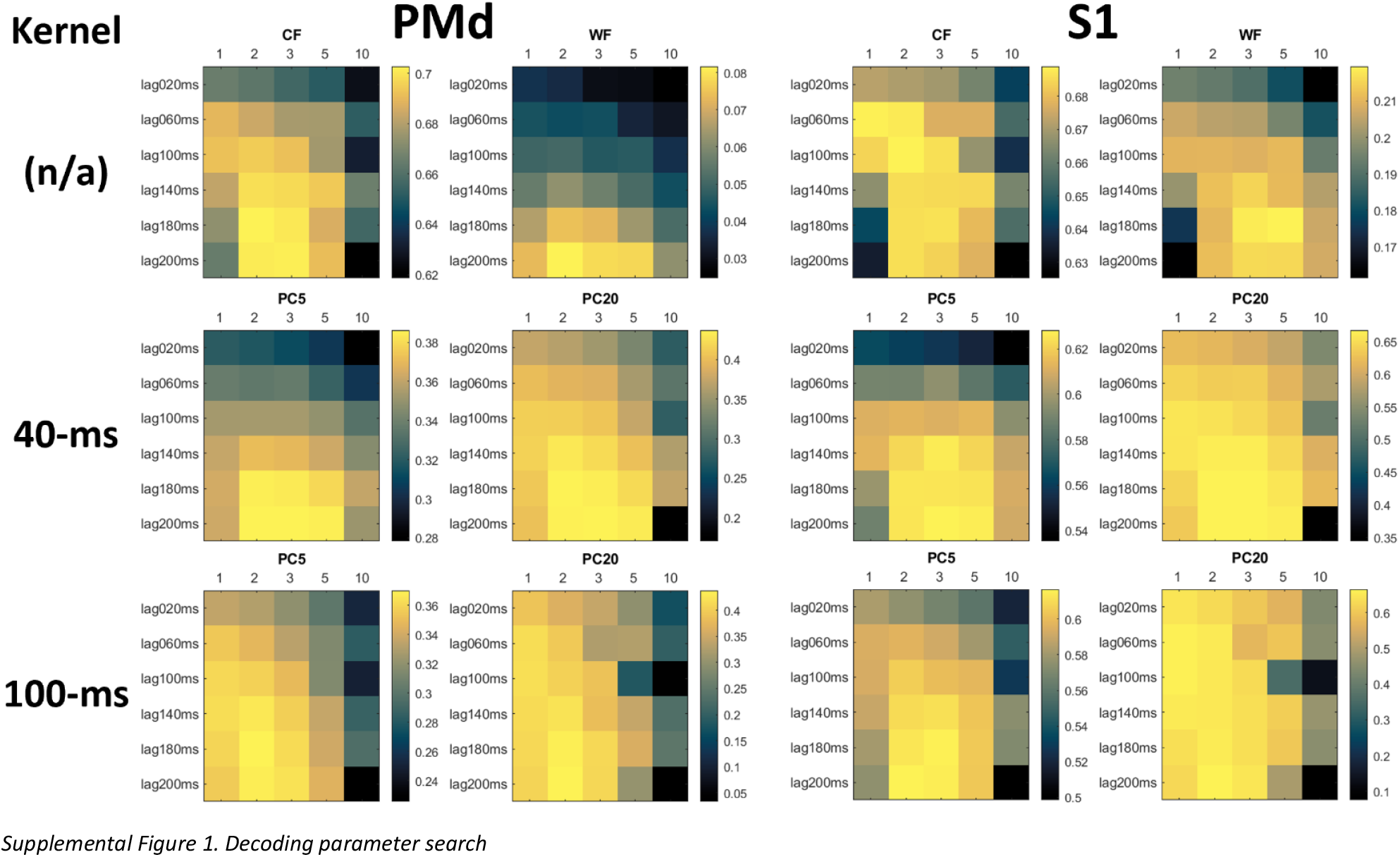
Decoding parameter search.

### SFig 2: Recovery: WFs

**Supplemental Figure 2.**
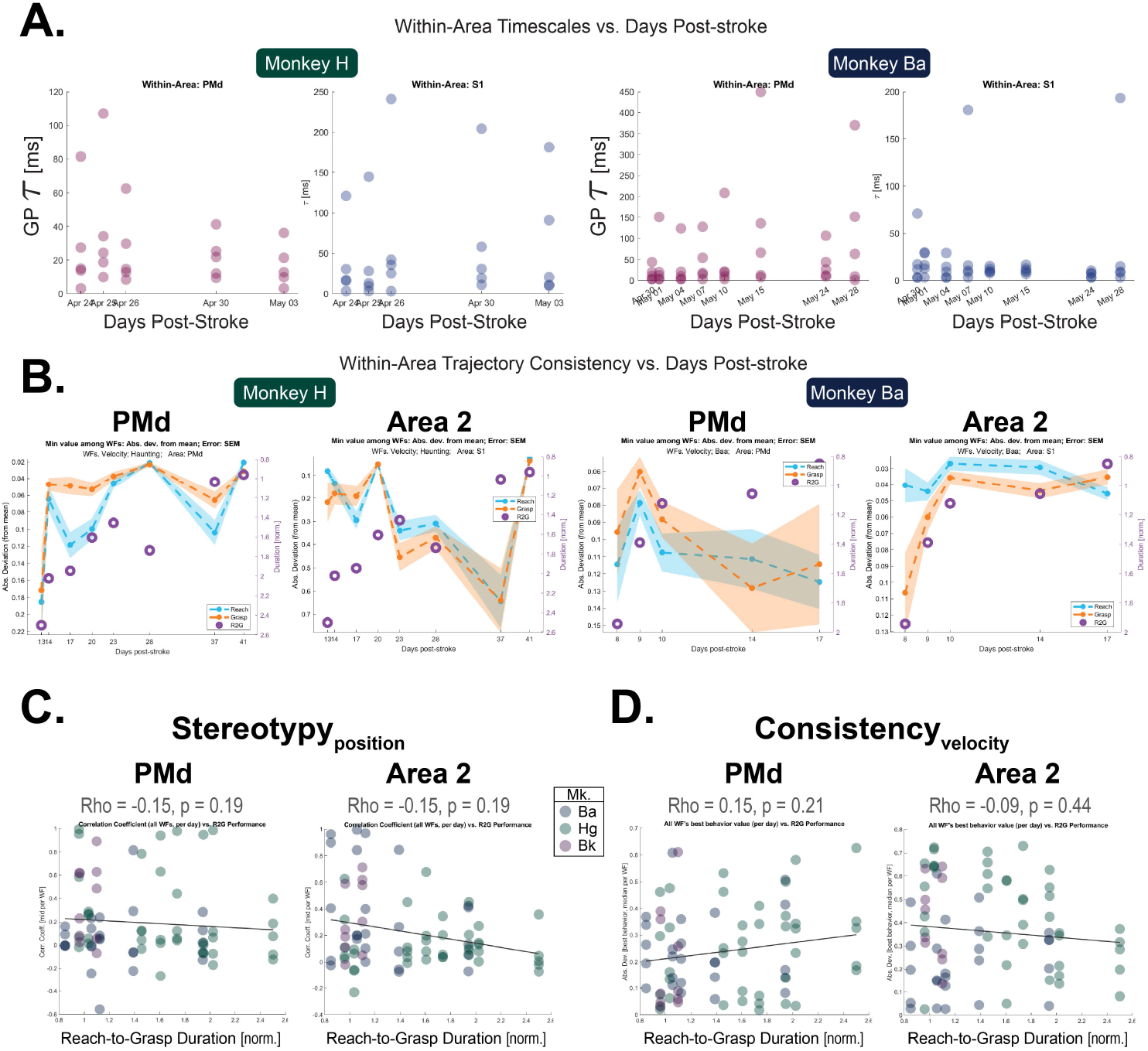
Within-area factors did not reliably track neural recovery. WF results for the same two animals that showed post-lesion behavioral deficits (n = 2) as shown in Figure 4 and Figure 5. All DLAG results in this figure are from models trained on low-SBP data. Values are separated per area (PMd and Area 2) for all tiles. (A)WF Gaussian process widths (τ) per session over time. Left two tiles: Mk. H; Right two tiles: Mk. Ba. (B)WF trajectory consistency (cyan and orange) and R2G performance (purple). Showing results for the most consistent WF for two R2G timepoint subsets: reach start (cyan) and grasp (orange); “grasp” is the mean of grasp start and grasp finish. Midlines: median per-trial consistency. Error: SEM. Normalized R2G duration per session (purple). (C)WF trajectory stereotypy_position_ vs. normalized R2G duration for all animals (n = 3). Per area, the three most stereotyped WFs for each session are included. Line shows linear model fit. (D)WF instantaneous consistency_velocity_ vs. normalized R2G duration for all animals (n = 3). Per area, the three most consistent WFs for each session are included. Line shows linear model fit.

